# Stathmin-2 Mediates Glucagon Secretion from Pancreatic α-Cells

**DOI:** 10.1101/832493

**Authors:** Farzad Asadi, Savita Dhanvantari

## Abstract

Inhibition of glucagon hypersecretion from pancreatic α-cells is an appealing strategy for the treatment of diabetes. Our hypothesis is that proteins that associate with glucagon within alpha cell secretory granules will regulate glucagon secretion, and may provide druggable targets for controlling abnormal glucagon secretion in diabetes. Recently, we identified a dynamic glucagon interactome within the secretory granules of the α cell line, αTC1-6, and showed that select proteins within the interactome could modulate glucagon secretion. In the present study, we show that one of these interactome proteins, the neuronal protein stathmin-2, is expressed in aTC1-6 cells and in mouse pancreatic alpha cells, and is a novel regulator of glucagon secretion. Stathmin-2 was co-secreted with glucagon in response to 55 mM K+, and immunofluorescence confocal microscopy showed co-localization of stathmin-2 with glucagon and the secretory granule markers chromogranin A and VAMP-2 in αTC1-6 cells. In mouse pancreatic islets, Stathmin-2 co-localized with glucagon, but not with insulin, indicating that it is an alpha cell protein. To show a function for stathmin-2 in regulating glucagon secretion, we showed that siRNA - mediated depletion of stathmin-2 in aTC1-6 cells caused glucagon secretion to become constitutive without any effect on proglucagon mRNA levels, while overexpression of stathmin-2 completely abolished both basal and K+-stimulated glucagon secretion. Overexpression of stathmin-2 increased the localization of glucagon into the endosomal-lysosomal compartment, while depletion of stathmin-2 reduced the endosomal localization of glucagon. Therefore, we describe stathmin-2 as having a novel role as an alpha cell secretory granule protein that modulates glucagon secretion via trafficking through the endosomal-lysosomal system. These findings describe a potential new pathway for the regulation of glucagon secretion, and may have implications for controlling glucagon hypersecretion in diabetes.

## Introduction

Hyperglucagonemia is a characteristic sign of diabetes, causing fasting hyperglycemia and glycemic volatility. Clinically, glycemic variability contributes to the development of diabetes complications (1). Persistent hyperglucagonemia may exacerbate abnormal glucose metabolism in patients with type 2 diabetes and lead to metabolic disturbances in obese and prediabetic individuals (2). Accordingly, controlling this excess glucagon secretion may be a potential therapeutic strategy for diabetes (3) so that glycemia and glucose metabolism may be better regulated. Such an approach has been suggested as a priority for the treatment of diabetes (1).

Combating hyperglucagonemia could be theoretically achieved by *i)* inhibition of glucagon action at target organs by blocking the glucagon receptor, or *ii)* inhibition of glucagon secretion from the pancreatic α cells. While in the short-term the former could be an effective therapeutic strategy, it can lead to α cell hyperplasia and hyperglucagonemia over a long-term period (4), along with a risk of hypoglycemia (5) and disturbances in lipid metabolism (6). Therefore, inhibiting glucagon secretion, rather than blocking the glucagon receptor, may be a more appropriate therapeutic approach for the treatment of hyperglucagonemia of diabetes (7).

It has been documented that suppression of glucagon secretion can be mediated at the systemic, paracrine or intrinsic level (8). As a systemic modulator, GLP-1 directly inhibits glucagon secretion from α cells by signaling through the alpha cell GLP-1R (9) (10) (11), or indirectly by increasing inter-islet somatostatin or insulin secretion (12) (9). Glucagon secretion is also suppressed by paracrine signaling through the insulin, somatostatin and GABA_A_ receptors on the α cell (13) (14) (12). At an intrinsic level, glucose directly or indirectly inhibits glucagon secretion from the α cell (15) (16) (17) (18) by altering downstream activities of Ca^2+^ channels, K_ATP_ channels (19), and trafficking of secretory granules (20).

We are pursuing the hypothesis that glucagon secretion can also be controlled by proteins within the secretory granule that associate with glucagon. By conducting secretory granule proteomics in αTC1-6 cells, we have recently described a dynamic glucagon interactome, and shown that components of this interactome can play a role in modulating glucagon secretion (21). Of these components, a protein of particular interest is Stathmin-2 (Stmn2 or SCG10), a member of the stathmin family of Golgi proteins (22), that may play a role in the regulation of neuroendocrine secretion (23). In the human islet, Stmn2 expression may be unique to alpha cells, as shown by genome-wide RNA-Seq analysis (24) and single cell transcriptomics (23), and alpha cell Stmn2 mRNA expression is differentially regulated in type 2 diabetes (25). These studies, together with our proteomics findings, led us to hypothesize that Stmn2 may function in α cells to modulate glucagon secretion. In the present study, we show that Stmn2 is co-localized with glucagon in secretory granules of αTC1-6 cells and controls glucagon secretion by trafficking through the endosomal/lysosomal system.

## Materials and Methods

### Cell culture

αTC1-6 cells (a kind gift from C. Bruce Verchere, University of British Columbia, Vancouver, BC, Canada) were cultured in regular DMEM medium containing 5.6 mM glucose (Cat# 12320032, Thermo Fisher Scientific) supplemented with 15% horse serum (Cat# 26050088, Thermo Fisher Scientific), 2.5% FBS (Cat# 16000044, Thermo Fisher Scientific), L-glutamine and sodium pyruvate. For secretion experiments, cells were plated in 6 –well plates and for all experiments, a low passage number (up to P6) was used. Twenty-four hours prior to secretion experiments, media were removed and replaced with DMEM without supplements. To evaluate (21) the regulated secretion of glucagon, cells were washed twice with HBSS, pre-incubated for 2h in DMEM without supplements, then incubated for 15 min with or without KCl (55 mM). These media were collected into tubes containing protease inhibitor (PMSF, 45 mM) and phosphatase inhibitors (sodium orthovanadate, 1mM; and sodium fluoride 5 mM) while tubes were kept on ice. After collection, media were centrifuged at 13000×g for 5 min at 4°C, and the supernatant was collected into new microfuge tubes and immediately kept at -80°C until analysis. After media were removed, cells were washed using ice-cold PBS (pH 7.4) and lysed in RIPA buffer (Cat# 89900, Thermo Fisher Scientific) containing abovementioned protease and phosphatase inhibitors. The lysed cells were centrifuged at 13000×g for 5 min at 4°C and the supernatant was kept at -80°C for protein assay.

### Gene construct and plasmid preparation

To generate the expression plasmid for Stmn2, the Kozak sequence (GCCACC), signal peptide sequence and coding sequence of mouse Stmn2 (https://www.uniprot.org/uniprot/P55821) were ligated into the NheI and ApaI restriction sites of pcDNA3.1(+) MAr. The construct was synthesized by GENEART GmbH, Life Technologies (GeneArt project 2018AAEGRC, Thermo Fisher Scientific). Then, Max Efficiency DH5α competent cells (Thermo Fisher Scientific, Cat# 18258012) were transformed according to the manufacturer’s protocol. Plasmids were then extracted and purified using the PureLink HiPure Plasmid Maxiprep Kit (Thermo Fisher Scientific, Cat# K210006) for downstream experiments. Correct assembly of the final construct was verified by gene sequencing at the London Regional Genomics Facility, Western University. For Stmn2 overexpression studies, αTC1-6 cells were transiently transfected by pcDNA3.1 (+) MAr-stmn2 construct or empty vector (negative control). All transfections were done using Lipofectamine 2000 (Cat#11668-027, Invitrogen). To monitor normal cell growth and morphology, cells were checked daily by the EVOS cell imaging system (Thermo Fisher Scientific).

### Gene silencing experiments

#### a) siRNA-mediated depletion of stathmin-2

Functional analysis of Stmn2 was done by gene silencing experiments. siRNAs targeting three regions within the Stmn2 mRNA (Cat# s73356, s73354, s73355) were chosen from pre-designed mouse siRNAs (Silencer siRNA, Thermo Fisher Scientific). The control group was treated with Mission siRNA Universal Negative Control # 1 (Cat# SIC001, Sigma-Aldrich). Gene silencing was done based on a previously published protocol (26) and as we have done previously with some modifications (21). Briefly, αTC1-6 cells were cultured in regular DMEM to 60% confluency. Media were removed and replaced with 2 mL Opti-MEM (Cat# 31985-070, Gibco) containing 50 nM pooled siRNAs with Lipofectamine 2000. After 8h, media were changed to regular DMEM without FBS and cultured for 72h. Then, media were refreshed and cells were cultured for 15 min in the absence of presence of 55 mM KCl as described above. Gene silencing was confirmed by analyzing mRNA expression levels of Stmn2. After removing media, cells were washed by cold PBS (pH 7.4) and total RNA was extracted (RNeasy extraction kit; Cat # 74104, Qiagen). cDNA synthesis was performed using the SuperScript III First Strand Synthesis Supermix for qRT-PCR (Cat # 11752050, Thermo Fisher Scientific), according to the supplier’s protocol. Real-time PCR was performed using Quant Studio Design and Analysis Real-Time PCR Detection System in conjunction with the Maxima SYBR Green qPCR Master Mix (Cat # K0221, Thermo Fisher Scientific) using specific primers for Stmn2: forward, 5′-GCAATGGCCTACAAGGAAAA-3′; reverse, 5′-GGTGGCTTCAAGATCAGCTC-3′; and β -Actin; forward, 5′-AGCCATGTACGTAGCCATCC-3′; reverse, 5′-CTCTCAGCTGTGGTGGTGAA-3′. Gene expression levels for stathmin-2 were normalized to that of β-Actin. The normalized level of transcripts in the depleted cells was shown relative to that of the non-targeting negative control. Relative expression levels were determined as percent of alterations compared to the control. Statistical analysis was performed using t-test at α = 0.05.

#### b) Proglucagon gene expression levels following Stmn2 depletion

After siRNA treatments, proglucagon gene expression levels were measured by real-time PCR as described above using proglucagon-specific primers (forward: 5′-CAGAGGAGAACCCCAGATCA-3′, reverse: 5′-TGACGTTTGGCAATGTTGTT-3′).

### Immunofluorescence confocal microscopy

#### αTC1-6 cells

To determine if Stmn2 and glucagon could be co-localized to the same intracellular compartments, we used immunofluorescence confocal microscopy. Wild type αTC1-6 cells were seeded on collagen (type I)-coated coverslips and grown in regular DMEM, then incubated in non-supplemented DMEM for 24h. At the end of incubation, cells were washed once with PBS, fixed in 2% paraformaldehyde (in PBS) for 30 min, permeabilized with 0.25% Triton X-100 (in PBS) for 5 min and washed with PBS. After 1h incubation with blocking buffer (10% goat serum in 1% BSA/PBS), coverslips were incubated with primary antibodies against glucagon (mouse anti-glucagon antibody, Cat # ab10988, Abcam; 1:1000), Stathmin-2 (goat anti-SCG10 antibody, Cat # ab115513, Abcam; 1:1000), secretory granule marker, chromogranin A (mouse anti-ChgA antibody, Cat# MAB319, Sigma; 1:1000), secretory granule marker, VAMP2 (rabbit anti-VAMP2, Cat# ab215721, Abcam; 1:1000), early endosome marker, EEA1 (rabbit antiEEA1, Cat # ab 2900, Abcam; 1:500) or the lysosomal marker, Lamp2A (rabbit anti-Lamp2A, Cat# ab18528, Abcam; 1:500) overnight. After washing with PBS, coverslips were incubated with the following secondary antibodies as appropriate: goat anti-mouse IgG Alexa Fluor 488 (Cat# A-11001, Molecular Probes; 1:500), goat anti-rabbit IgG Alexa Fluor 594 (Cat# A11037, Invitrogen; 1:500) or donkey anti-goat IgG Alexa Fluor 555 (Cat# ab150130, Abcam; 1:500) Invitrogen) for 2h in the dark at room temperature. Then, coverslips were washed with PBS and mounted on glass slides using DAPI containing ProLong antifade mountant (Cat # P36935, Molecular Probes) for image analysis by confocal immunofluorescence microscopy (Nikon A1R, Mississauga, Canada). Coverslip preparation was done at four different times with freshly thawed cells.

#### Mouse pancreatic islets

All mice were treated in accordance with the guidelines set out by the Animal Use Subcommittee of the Canadian Council on Animal Care at Western University based on the approved Animal Use Protocol AUP 2012-020. Six to 8-week old male C57BL/6 mice (n=7) were sacrificed by cervical dislocation under anesthesia with inhalant isoflurane. Pancreata were collected and fixed in 10% buffered formalin for 3 days and treated with 70% ethanol for one day before paraffin embedding at Molecular Pathology Department, Robarts Research Institute, Western University. The paraffin-embedded blocks were longitudinally sectioned in 5 µm slices and fixed onto glass micrscope slides. The samples were de-paraffinized by graded washes using xylene, ethanol and PBS. Background Sniper (Cat# BS966H, Biocare Medical) was used to reduce non-specific background staining. Samples were incubated with primary antibodies against glucagon (1:500), Stmn2 (1:250), insulin (Cat# ab7842, Abcam; 1:250) and TGN46 (Cat# ab16059, Abcam; 1:200) and followed by secondary antibodies of goat anti-mouse IgG Alexa Fluor 488 (1:500), donkey anti-goat IgG Alexa Fluor 555 (1:500), and goat anti-guinea pig IgG Alexa Fluor 647 (Cat# A21450, Invitrogen; 1:500). Nuclei were stained with DAPI (1:1000), and tissues were mounted in Prolong antifade mountant (Cat# P36982, Thermo Fisher Scientific). As a background control for Stmn2, islet staining for Stmn2 was done using only the secondary antibody.

#### Image acquisition

Images were acquired through a Nikon A1R Confocal microscope with a ×60 NA plan-Apochromat oil differential interference contrast objective and NIS-Elements software (Nikon, Mississauga, Canada). High-resolution images were acquired by selecting Nyquist XY scan area, 1024×1024 pixel size scanning of the selected area and 2D-deconvolution of the captured images.

#### Image Analysis

For cell image analysis, we prepared three coverslips for each group. Image analysis was performed by NIS-Elements software (Nikon, Mississauga, Canada), using the co-localization option and Pearson’s correlation coefficient (PCC). Regions of interest (ROI) were manually drawn around distinct single or multicellular bodies, and merged values of glucagon and Stmn2 were taken for analysis. Colocalization of the pixels from each pseudo-colored image was used to calculate Pearson’s correlation coefficient, as we described previously (27) (21).

For mouse pancreatic islets, images were captured using four channels of green (glucagon), red (Stmn2), purple (insulin) and blue (nucleus; DAPI). To calculate the extent of co-localization between glucagon and stathmin-2 (glucagon^+^, Stmn2^+^), images of 15 islets per pancreas were captured and analysed by Pearson’s correlation coefficient (PCC). To this end, we manually drew ROIs around each islet and then defined PCC values for colocalization between Stmn2 and glucagon or insulin using the colocalization option of the NIS-Elements software. To predict expression levels of Stmn2 in α or β-cells of the pancreatic islets we have performed binary analysis using M-Thresholding algorithm of NIS-Elements software, followed by regression analysis of Stmn2 versus glucagon or insulin using GraphPad Prism 7.

### Measurement of glucagon and stathmin-2

Glucagon levels in the media were determined by ELISA (Cat # EHGCG, Thermo Fisher Scientific) according to the manufacturer’s instructions. Stmn2 levels in the media were measured using mouse stathmin-2 ELISA kit (Cat# MBS7223765, MyBioSource) according to the manufacturer’s instruction. For each measurement, the values were compared between groups by t-test and among groups by 1-Way ANOVA (α = 0.05). Cell protein levels were determined using BCA assay and used for normalization of the glucagon or Stmn2 levels.

## Results

### Stmn2 co-localizes with glucagon and secretory granule markers in αTC1-6 cells

Immunostaining of glucagon and Stmn2 in αTC1-6 cells revealed significant co-localization, as shown in Figure 1A and by a positive Pearson’s correlation coefficient (0.74±0.05) between endogenously expressed glucagon and Stmn2. Linear regression of binary intensities showed a sensitive and significant relationship between colocalization of glucagon and Stmn2 (Figure 1B). Both glucagon and stathmin-2 were co-secreted in response to 55 mM K^+^, with corresponding decreases in cell contents (Figures 1C and D). To further confirm the presence of stathmin-2 in secretory granules, aTC1-6 cells were immunostained for stathmin-2 and the secretory granule markers chromogranin A (Figure 1E) and VAMP2 (Figure 1F). There was moderate colocalization (28) between glucagon and chromogranin A (PCC 0.58 ± 0.07) or VAMP2 (PCC 0.56 ± 0.09) (Figure 1G), indicating that Stmn2 is partially localized to the secretory granule compartment in αTC1-6 cells.

**Figure 1:**
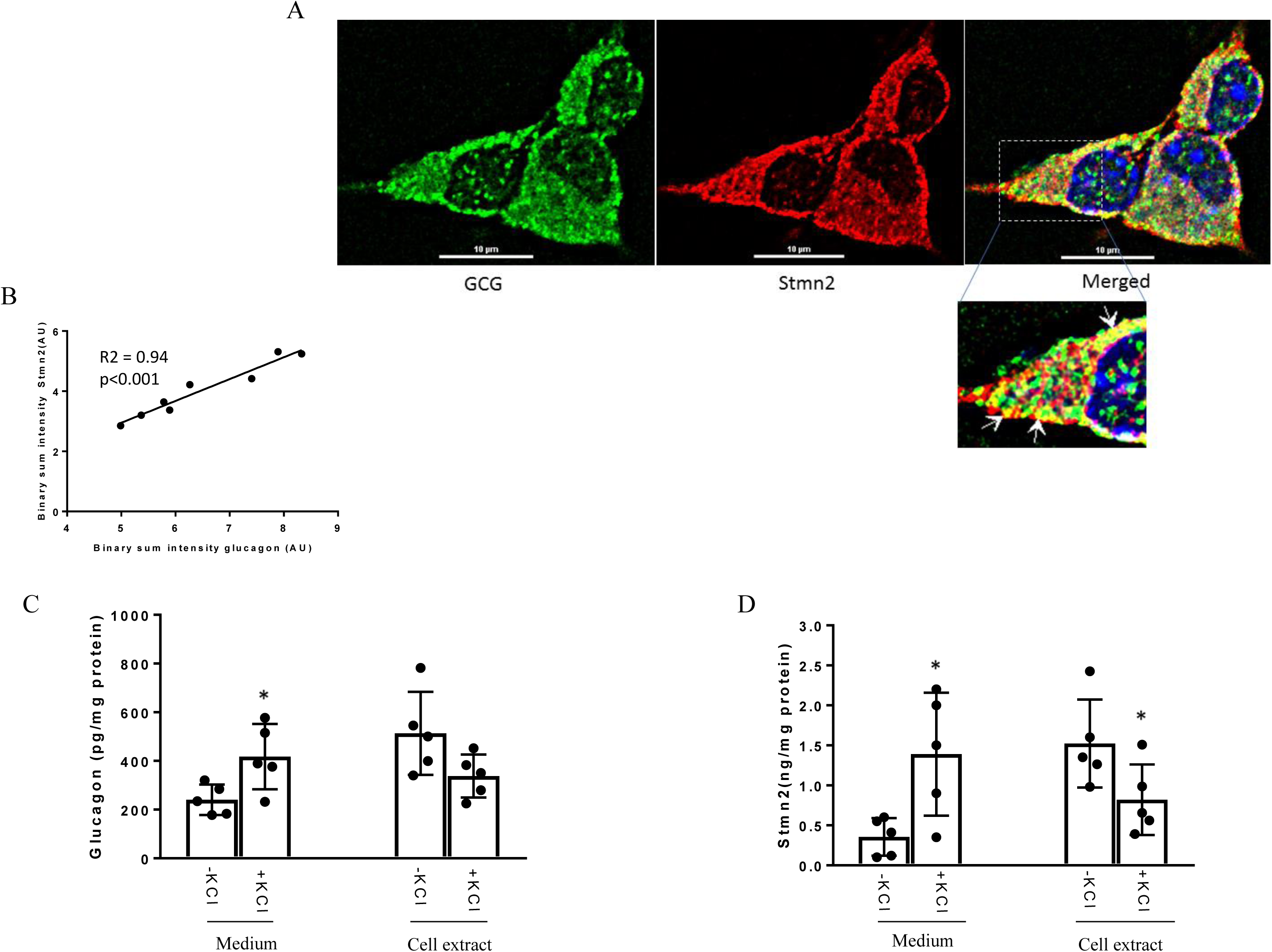
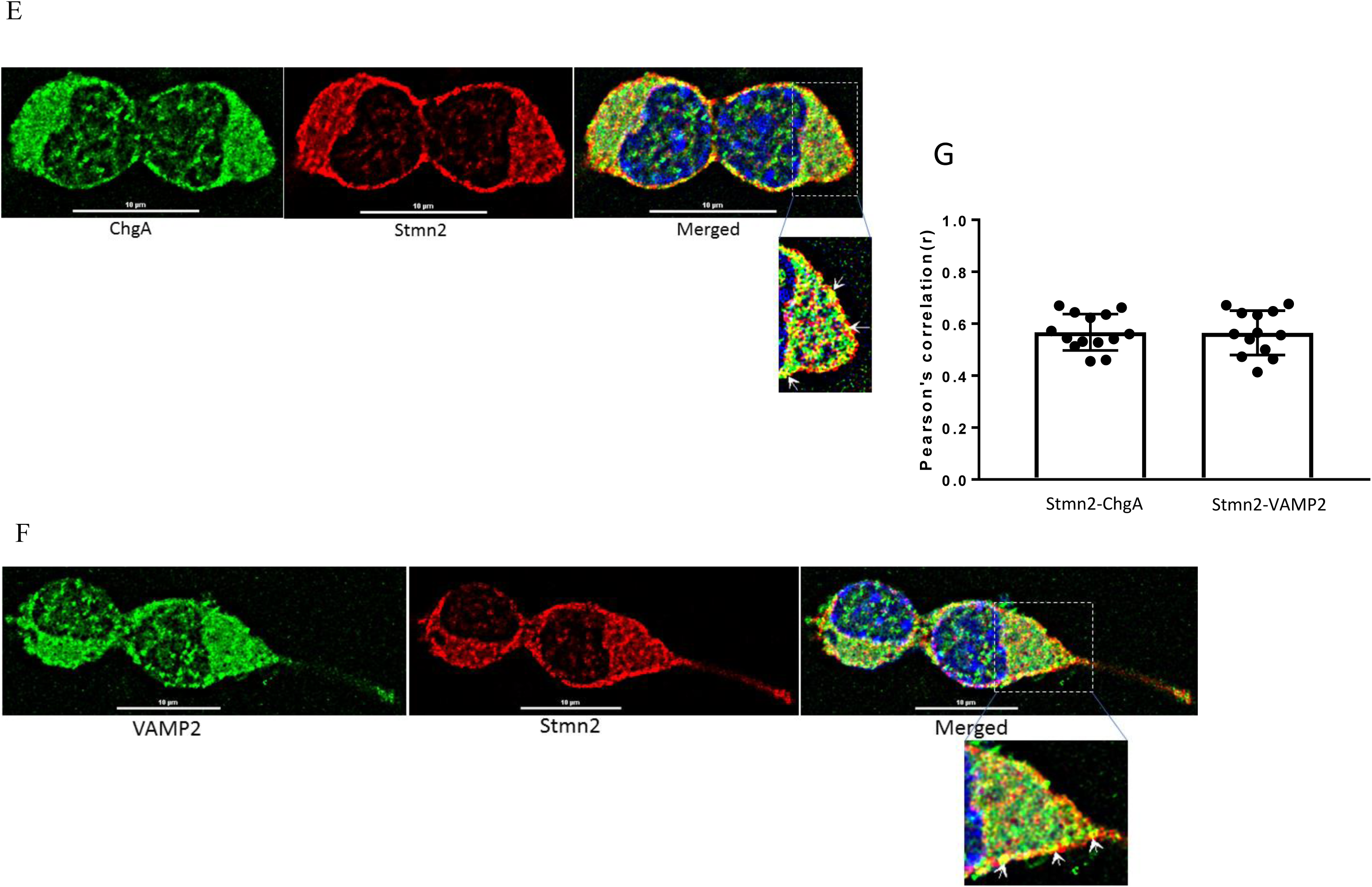
Stathmin-2 localizes to secretory granules in αTC1-6 cells. αTC1-6 cells were immunostained using primary antibodies against glucagon (GCG, green) and stathmin-2 (Stmn2, red). DAPI (blue) indicates the nucleus in the merged image. Resolution of the images was extended by applying Nyquist XY scan and then 2D- Deconvolution in NIS Elements image analysis software. Images are representative of four biological replicates with 3 technical replicates each. **(A)** Areas of yellow in the merged image show colocalization of glucagon and Stmn2. (B) Linear regression analysis of binary intensities of glucagon and Stmn2 predicts a significant (p<0.001) correlation. Each value represents mean intensities of 5-7 cells. The secretion of both glucagon (C) and Stmn2 (D) was significantly increased after treatment with 55 mM KCl for 15 min. Values are expressed as mean ± SEM (n=5). *p<0.05. Stmn2 colocalizes with the secretory granule proteins ChgA (E) and VAMP2 (F), as indicated by yellow punctate staining. (G) The extent of colocalization was analysed by Pearson correlation coefficient for Stmn2 with ChgA or with VAMP2.

### Stathmin-2 localizes to α cells in mouse pancreatic islets

Immunostaining of mouse pancreatic islets showed a pattern of Stmn2 immunofluorescence similar to that of glucagon, and not insulin (Figure 2A). Analysis by Pearson correlation showed a strong colocalization between glucagon and Stmn2 in the islets (PCC = 0.77± 0.02), but not between insulin and Stmn2 (Figure 2B). Linear regression analysis of the binary intensities revealed a very strong and significant relationship between Stmn2 and glucagon immunofluorescence (Figure 2C), while there was no significant relationship between Stmn2 and insulin (Figure 2D). In addition, Stmn2 showed a strong relationship with the *trans* Golgi marker, TGN46 (PCC = 0.72 ± 0.09). These results indicate that Stmn2 is localized to the secretory pathway of α cells in mouse pancreatic islets.

**Figure 2:**
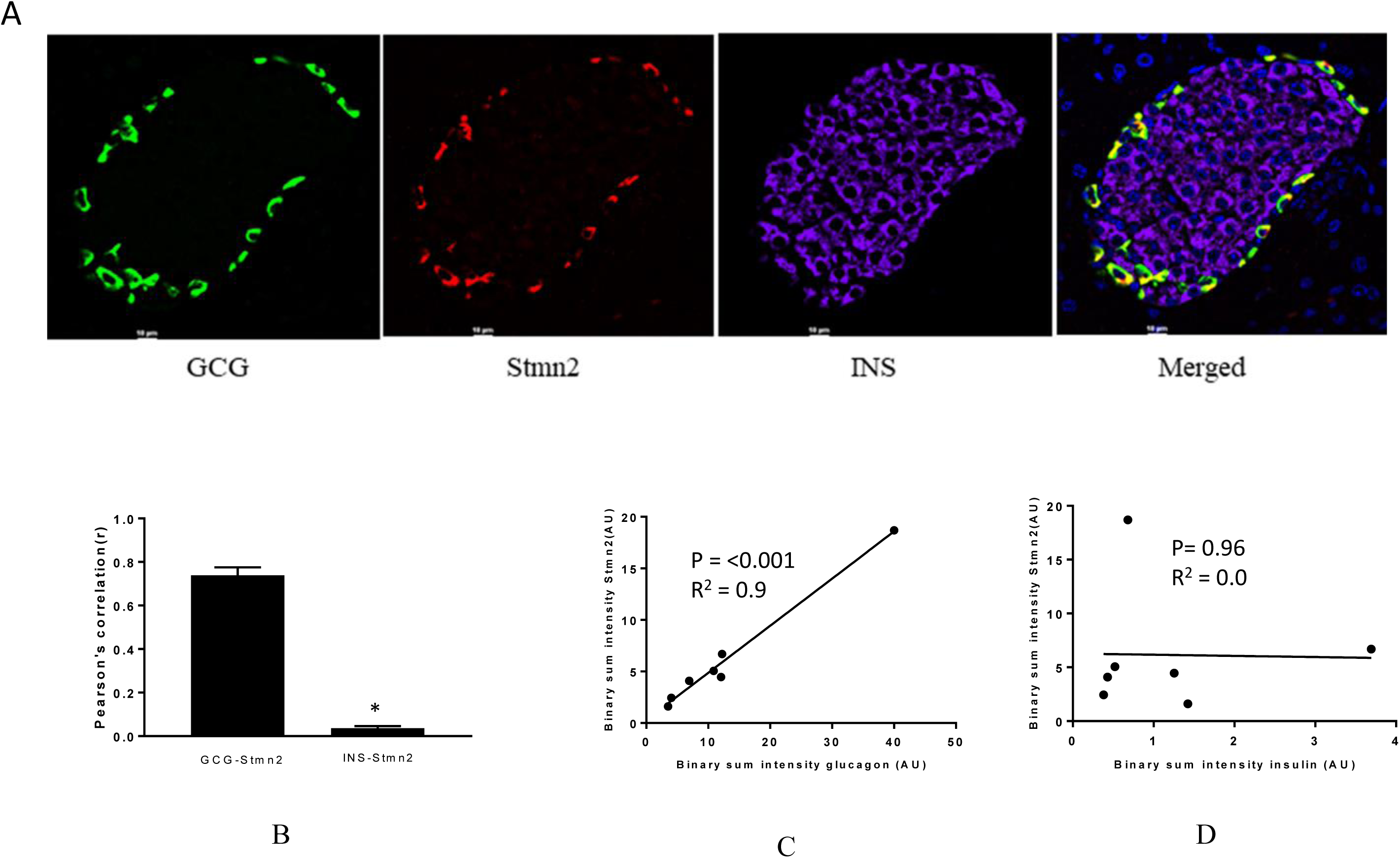
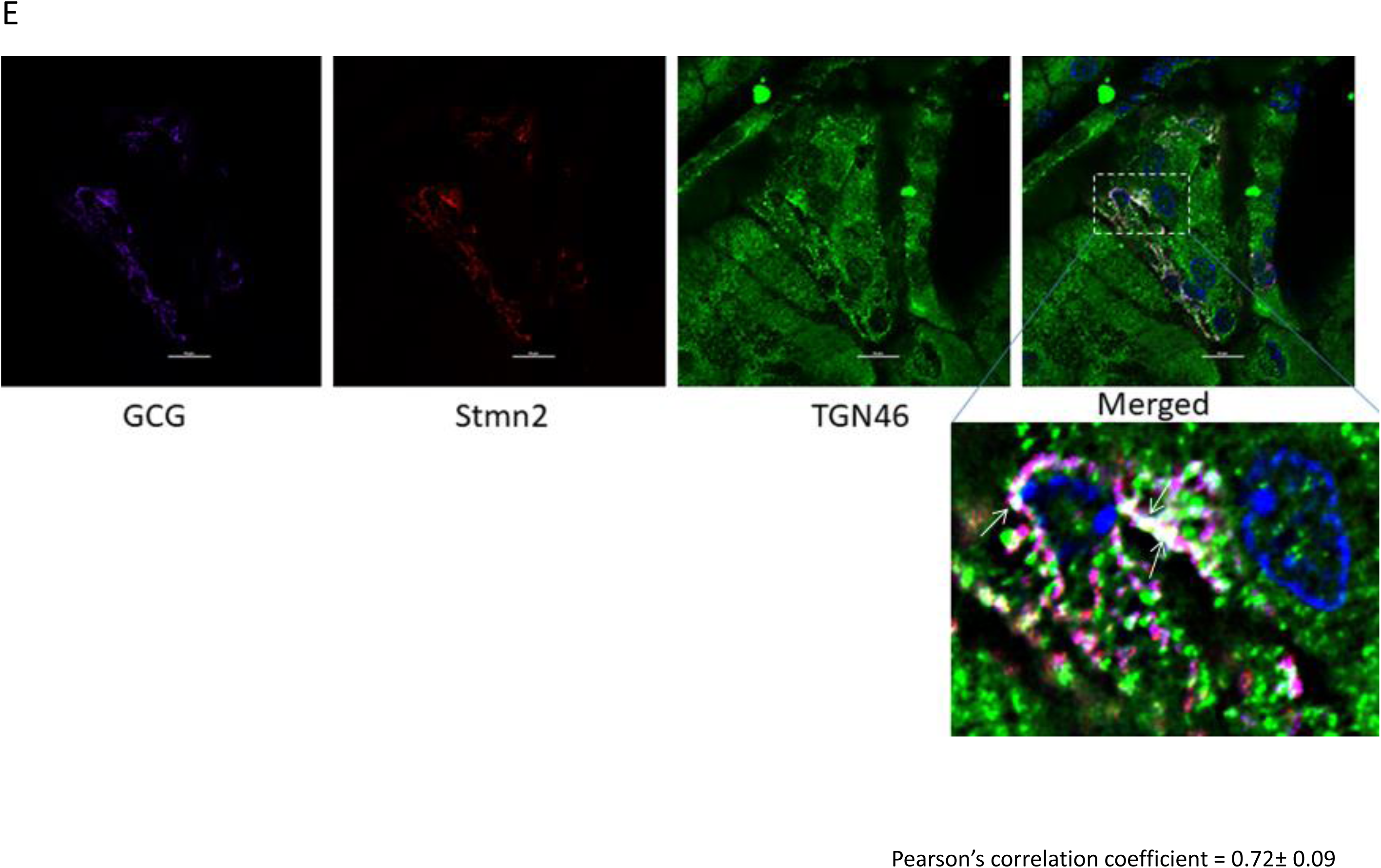
Stathmin-2 is present in α-cells, but not β cells, in murine pancreatic islets. Pancreata of C57BL/6 mice (n=7; 5µm sections) were immunostained for against glucagon (GCG), stathmin-2 (Stmn2) and insulin (INS). Images were acquired and analysed for co-localization as described in Figure 1. **(A)** Both glucagon and Stmn2 localize to the mantle of the islets, and areas of yellow in the merged image demonstrate dual positive alpha cells (glucagon^+^ and Stmn2^+^). **(B)** Pearson’s correlation coefficient for colocalization of Stmn2 and glucagon or insulin. (C) Linear regression analysis predicts a strong positive correlation between the binary intensities of glucagon and Stmn2 (p<0.001). D) There is no correlation between the binary intensities of insulin and Stmn2. (E) Colocalization of Stmn2 and the trans-Golgi marker TGN46 in murine pancreatic islets. Areas of white (arrows in the magnified panel) indicate co-localization of glucagon, Stmn2 and TGN46.

### Effects of depletion and overexpression of Stmn2 on glucagon secretion

In order to determine if Stmn2 had any functional effects in α-cells, we manipulated levels of Stmn2 and measured K+-stimulated glucagon secretion. Following siRNA-mediated knockdown of Stmn2 in αTC1-6 cells, basal secretion of glucagon was increased 6-fold, and was not significantly different from K+-stimulated secretion (Figure 3A), indicating increased constitutive secretion. Efficacy of siRNA-mediated depletion of Stmn2 was shown by a significant reduction (p<0.01) in the expression of Stmn2 mRNA levels (Figure 3B). As well, silencing of Stmn2 did not affect proglucagon gene expression levels (Figure 3C), indicating that the effects of Stmn2 depletion were on secretion alone. Conversely, overexpression of Stmn2 dramatically reduced both basal (∼6 times) and stimulated (∼11 times) glucagon secretion compared to the corresponding control groups (Figure 3D; p<0.001). These findings suggest that Stmn2 levels may control the regulated secretion of glucagon from αTC1-6 cells.

**Figure 3:**
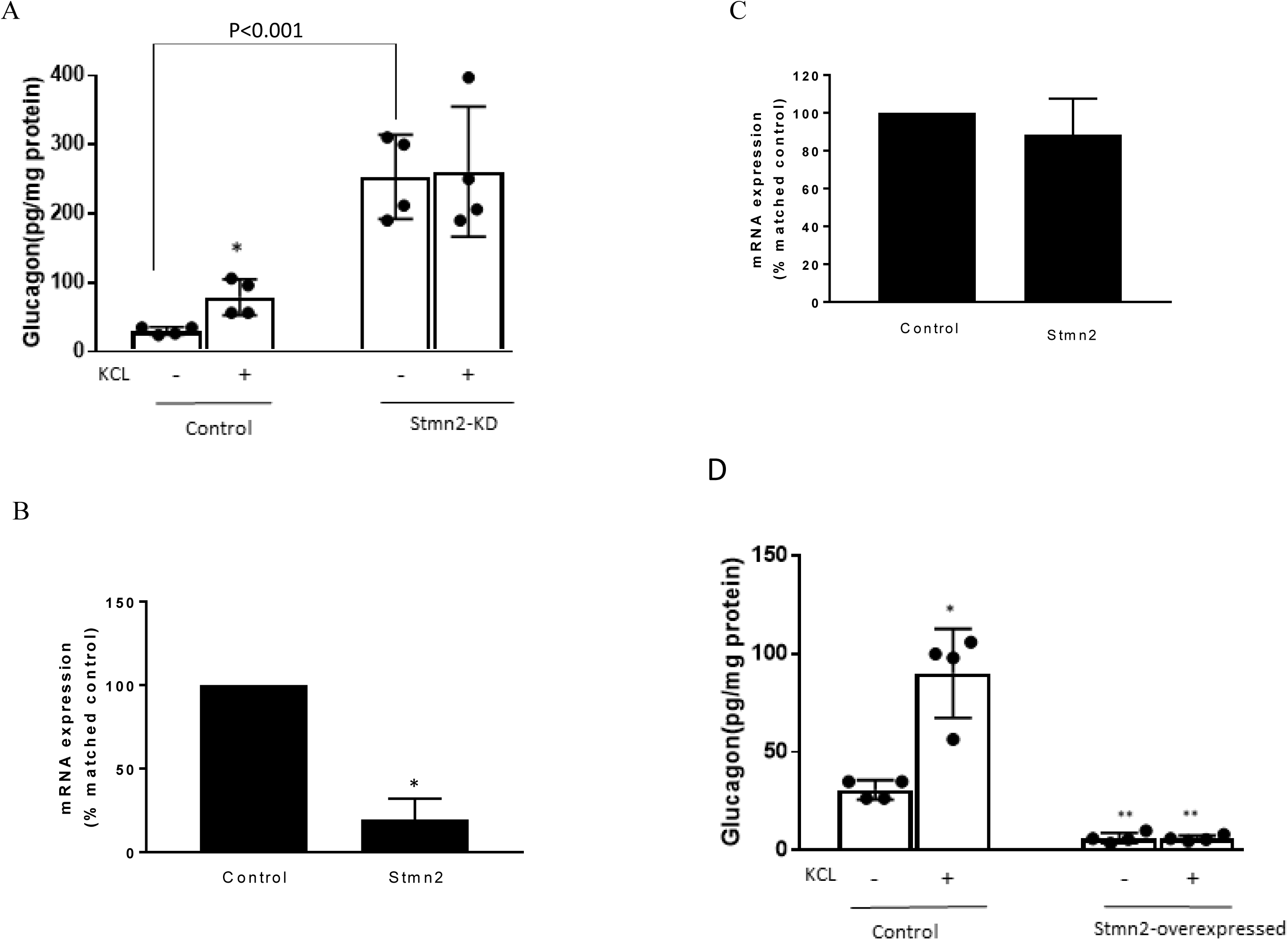
Silencing Stathmin-2 increased glucagon secretion and overexpression of stathmin-2 suppressed glucagon secretion in αTC1-6 cells. Wild type (wt; control) and stathmin-2 depleted (Stmn2-KD) αTC1-6 cells were preincubated 2h in serum free medium and then incubated with or without KCl (55 mM) for 15 min. (A) Glucagon secretion is significantly stimulated by KCL in wt cells, while in Stmn-KD cells, basal glucagon secretion is increased and does not respond to KCl. * p<0.01 compared to basal secretion in wt cells. (B) Stmn2 mRNA levels are decreased by about 70% after siRNA-mediated depletion in αTC1-6 cells. Values are means ± SEM (n=5), * p<0.01. **(C)** Proglucagon mRNA levels are not affected by siRNA-mediated depletion of stathmin-2. (D) Glucagon secretion is inhibited by overexpression of Stmn2. αTC1-6 cells were transfected with pcDNA3.1 (+) MAr-stmn2 construct or empty vector (negative control). Both basal (p<0.001) and K+-stimulated (p<0.001) glucagon secretion were inhibited by overexpression of Stmn2. Values are means ± SEM (n=4).

### Stmn2 directs glucagon into early endosomes

In wild type αTC1-6 cells, there was weak co-localization between glucagon and the early endosome marker EEA1 (Figure 4A) (PCC = 0.15±0.02). When Stmn2 was overexpressed (Figure 4B), the extent of colocalization between glucagon and EEA1 increased markedly (PCC = 0.53±0.08). Depletion of Stmn2 (Figure 4C) drastically reduced the extent of colocalization (PCC = 0.05) between glucagon and EEA1. Pearson’s correlation coefficient of co-localization between glucagon and Stmn2 showed a significant increase when Stmn2 was overexpressed (p<0.001) and a significant reduction when Stmn2 was knocked down (p<0.05) compared to the control (Figure 4D). These findings suggest that Stmn2 plays a role in directing glucagon towards early endosomes.

**Figure 4:**
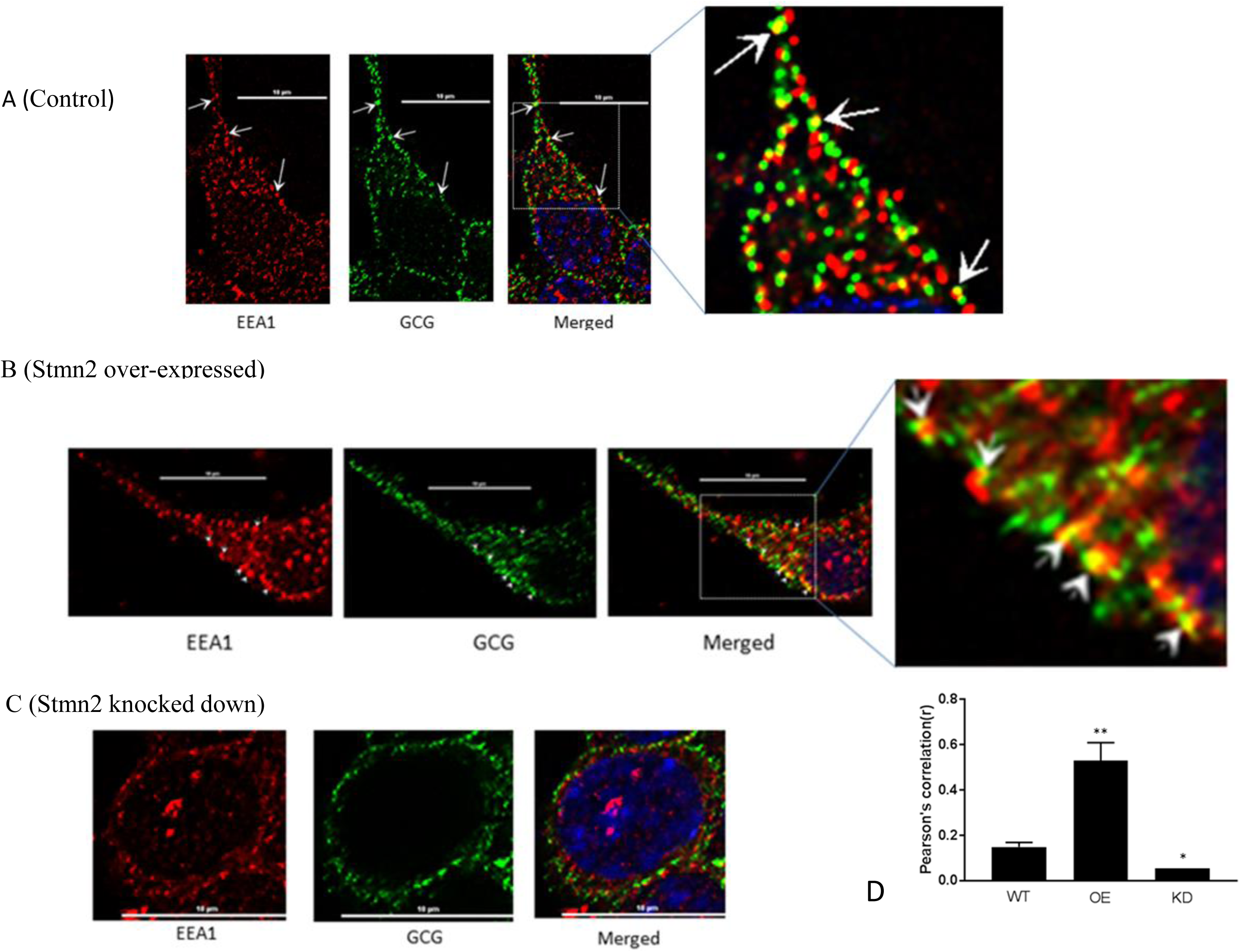
Stathmin-2 modulates glucagon trafficking through early endosomes. After transfection with either empty vector (A), vector encoding Stmn2 (B) or siRNAs against Stmn2 (C), αTC1-6 cells were immunostained using primary antibodies against glucagon and the early endosome marker EEA1. Images were acquired and analysed for co-localization as described in Figure 1. Colocalization of glucagon and EEA1 (yellow puncta) are indicated by arrows in wt cells (A) or arrowheads in cells overexpressing Stmn2 (B). (**D**) Level of colocalization between glucagon and EEA1 was determined by Pearson’s correlation coefficient in wt cells, cells in which Stmn2 was overexpressed (OE) and in which Stmn2 was knocked down by siRNA (KD). Values were expressed as mean ± SEM (n=5) and compared by 1-Way ANOVA. *p<0.05; **<0.001 compared to wt.

### Stmn2 overexpression increases glucagon presence in the late endosome/lysosome compartment

Similar to our findings in early endosomes, there was a weak correlation between glucagon and the late endosome-lysosome marker, Lamp2A (PCC = 0.2±0.02) in wild type αTC1-6 cells (Figures 5A, D). Following overexpression of Stmn2, the levels of colocalization between glucagon and Lamp2A were significantly increased (PCC=0.89±0.05, p<0.001) (Figures 5B, D). Depletion of Stmn2 significantly reduced the extent of colocalization between glucagon and Lamp2A compared to wild type cells (PCC = 0.001, p<0.01) (Figures 5C, D). Interestingly, the signal intensity of the endo-lysosomal marker, Lamp2A, was significantly increased (p < 0.01) upon overexpression of Stmn2, but not upon depletion of Stmn2 (Figure 5E).

**Figure 5:**
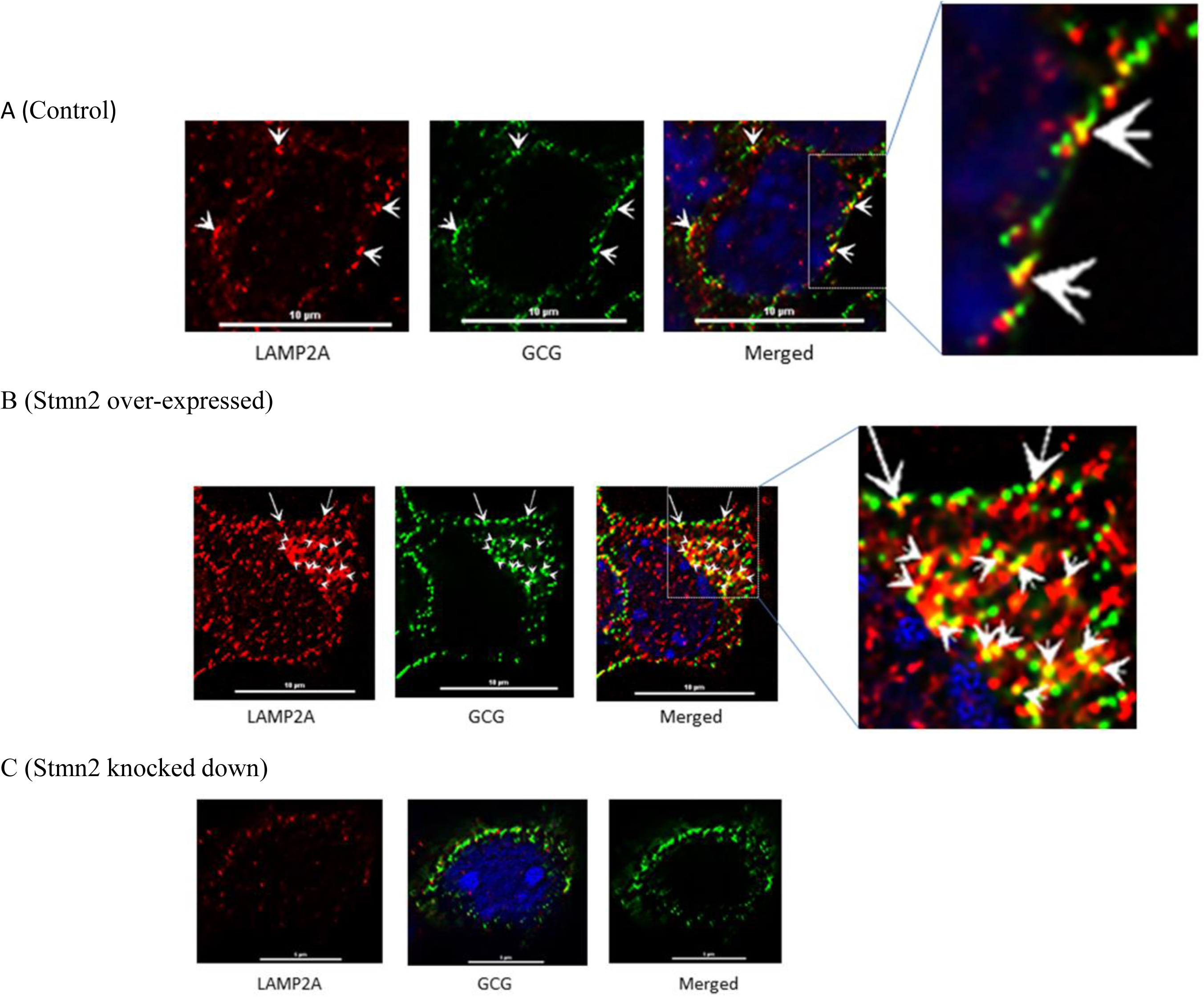
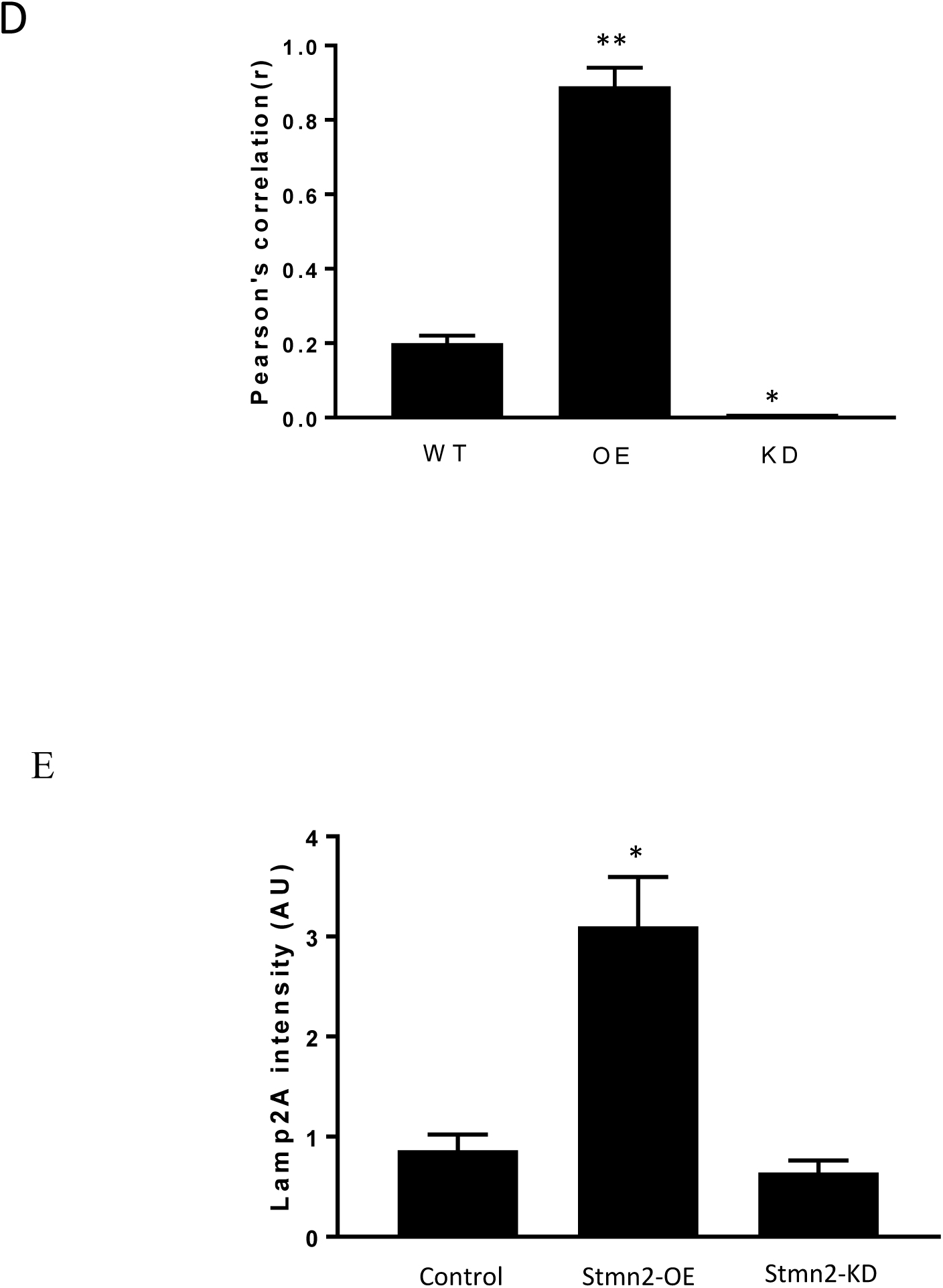
Overexpression of Stathmin-2 increases the presence of glucagon in late endosomes. After transfection with either empty vector (A), vector encoding Stmn2 (B) or siRNAs against Stmn2 (C), αTC1-6 cells were immunostained using primary antibodies against glucagon and the late endosome-lysosome marker LAMP2A. Images were acquired and analysed for co-localization as described in Figure 1.(A) In wt cells, glucagon and LAMP2 show some co-localization at the plasma membrane (arrowheads). (B) In cells overexpressing Stmn2, there is colocalization of glucagon and LAMP2A in the cell body (arrowheads) and at the plasma membrane (arrows). (C) In cells in which Stmn2 levels are depleted, there is almost no detectable co-localization of glucagon and LAMP2A. (**D**) Level of colocalization between glucagon and LAMP2A was determined by Pearson’s correlation coefficient in wt cells, cells in which Stmn2 was overexpressed (OE) and in which Stmn2 was knocked down by siRNA (KD). **(E)** Fluorescence intensity of Lamp2A in wt cells, and following overexpression (Stmn2-OE) or knockdown (Stmn2-KD) of Stmn2. Values are expressed as mean ± SD and compared by 1-Way ANOVA. *p<0.01; **<0.001 compared to wt.

## Discussion

Our overall hypothesis states that glucagon secretion from α cells is mediated by proteins that are associated with glucagon within secretory granules. To this end, we have shown that a neuronal protein, Stmn2, can be localized to the secretory granules of αTC1-6 cells. We validated this association in mouse pancreatic islets, and through silencing and overexpression experiments, we showed that Stmn2 can play a role in glucagon secretion by trafficking through the endosomal-lysosomal pathway.

We have previously identified Stmn2 as part of a network of proteins that associate with glucagon within the secretory granules of αTC1-6 cells (21). Data in the current study show that Stmn2 is localized to the secretory granule and Golgi compartments in aTC1-6 cells and mouse pancreatic alpha cells, respectively. Stathmin 2 is part of a family of neuronal phosphoproteins that associates with intracellular membranes, notably the Golgi and vesicle transporters, in neurons (29). Although Stmn2 has been identified as a neuron-specific protein that functions in differentiation and development, its presence in pancreatic alpha cells is not surprising, as several types of neuronal proteins, such as SNARE proteins, neurotransmitters and granins, are also expressed in endocrine cells (30) (31) (32). The expression of neuronal proteins such as Stmn2 in alpha cells may be due to the absence of the transcriptional repressor RE-1 silencing transcription factor (REST) in mature endocrine cells (33) (34). The absence of Stmn2 in mouse pancreatic beta cells suggests that transcriptional silencing programs may operate in a cell-specific manner.

Our results indicate that Stmn2 is localized largely to punctate structures within aTC1-6 cells, notably at the plasma membrane. Its colocalization with glucagon, ChGA and VAMP2 at the plasma membrane suggests that it is efficiently sorted from the Golgi to plasma membrane-associated secretory granules. The trafficking of Stmn2 to secretory granules in aTC1-6 cells may occur through specific molecular domains within its sequence. The subcellular trafficking pattern of Stmn2 in neurons and neuroendocrine cells is determined by its N-terminal extension, which contains a Golgi localization domain and a membrane anchoring domain that contains two conserved Cys residues as sites for palmitoylation (29). This lipid modification occurs in the Golgi (35) and is sufficient and necessary for the association of Stmn2 with Golgi membranes, and its sorting to post-Golgi vesicles (35) (36). While we have not directly shown that palmitoylation of these Cys residues is required for localization in secretory granules in aTC1-6 cells, it is likely that this is a conserved sorting mechanism in neuroendocrine cells.

Consistent with its localization within secretory granules, Stmn2 is co-secreted with glucagon in response to K+. While Stmn2 is a membrane–bound protein, it is also present in normal human blood as determined by plasma proteomics (37) and therefore can be also secreted. There is some evidence of a lower molecular weight form of Stmn2 that may correspond to a cleaved, soluble form (36) (38). In our model, it may be that a portion of Stmn2 becomes cleaved within the secretory granule, thus becoming part of the soluble cargo that is released.

By manipulating the expression of Stmn2, we were able to demonstrate a role in the control of glucagon secretion. There is precedence for a role for Stmn2 in the regulation of neuroendocrine secretion that has some interesting parallels with our results. While we showed that silencing of Stmn2 increased the constitutive secretion of glucagon, one study showed that silencing Stmn2 in PC12 cells decreased both basal and stimulated secretion of chromogranin A (23). In that study, it was shown that Stmn2 interacted directly with ChgA and its depletion reduced the buoyant density of chromaffin granules, suggesting that Stmn2 may participate in secretory granule formation, perhaps in partnership with ChgA (35)and thus promote regulated secretion. Although we could not demonstrate a direct interaction between Stmn2 and glucagon (data not shown), our results align with the idea that Stmn2 may be a sorting partner for glucagon, perhaps by interacting with other granule proteins such as ChgA or carboxypeptidase E (27), in the regulated secretory pathway of alpha cells.

A further clue to the mechanism of Stmn2 in the regulation of glucagon secretion was provided by overexpression experiments, in which we showed an almost complete shutdown of glucagon secretion and an increase in the localization of glucagon in early endosomes and lysosomes. These results suggest a mechanism whereby an increase in Stmn2 in the alpha cell inhibits glucagon secretion by targeting glucagon secretory granules for degradation in the endosome-lysosome pathway, perhaps in a manner similar to that of insulin secretory granules in Type 2 diabetes (39). Very recently, it has been shown that insulin secretion can be inhibited by the targeting of proinsulin for lysosomal degradation by Rab7-interacting lysosomal protein (RILP) (40). The proposed mechanism of action was through interactions between RILP and an insulin secretory granule membrane protein, Rab26. It is tempting to speculate that Stmn2 could have a similar mechanism of action in alpha cells. The increase in LAMP2 fluorescence intensity upon Stmn2 overexpression further suggests a role for Stmn2 in modulating glucagon secretion through increased lysosome function or biogenesis. We are currently investigating this mechanism of control of glucagon secretion in mouse islets.

In conclusion, we propose that Stmn2, a protein that is associated with glucagon in secretory granules, modulates glucagon secretion in alpha cells by playing a role in its intracellular trafficking. Under conditions that decrease Stmn2 levels, constitutive secretion of glucagon is increased; and under conditions that increase levels of Stmn2, glucagon is targeted to the endolysosomal system, presumably for degradation (Figure 6). Our findings represent a potentially novel intracellular pathway for the control of glucagon secretion, and may lead to new mechanistic insights in the dysregulation of glucagon secretion in diabetes.

**Figure 6:**
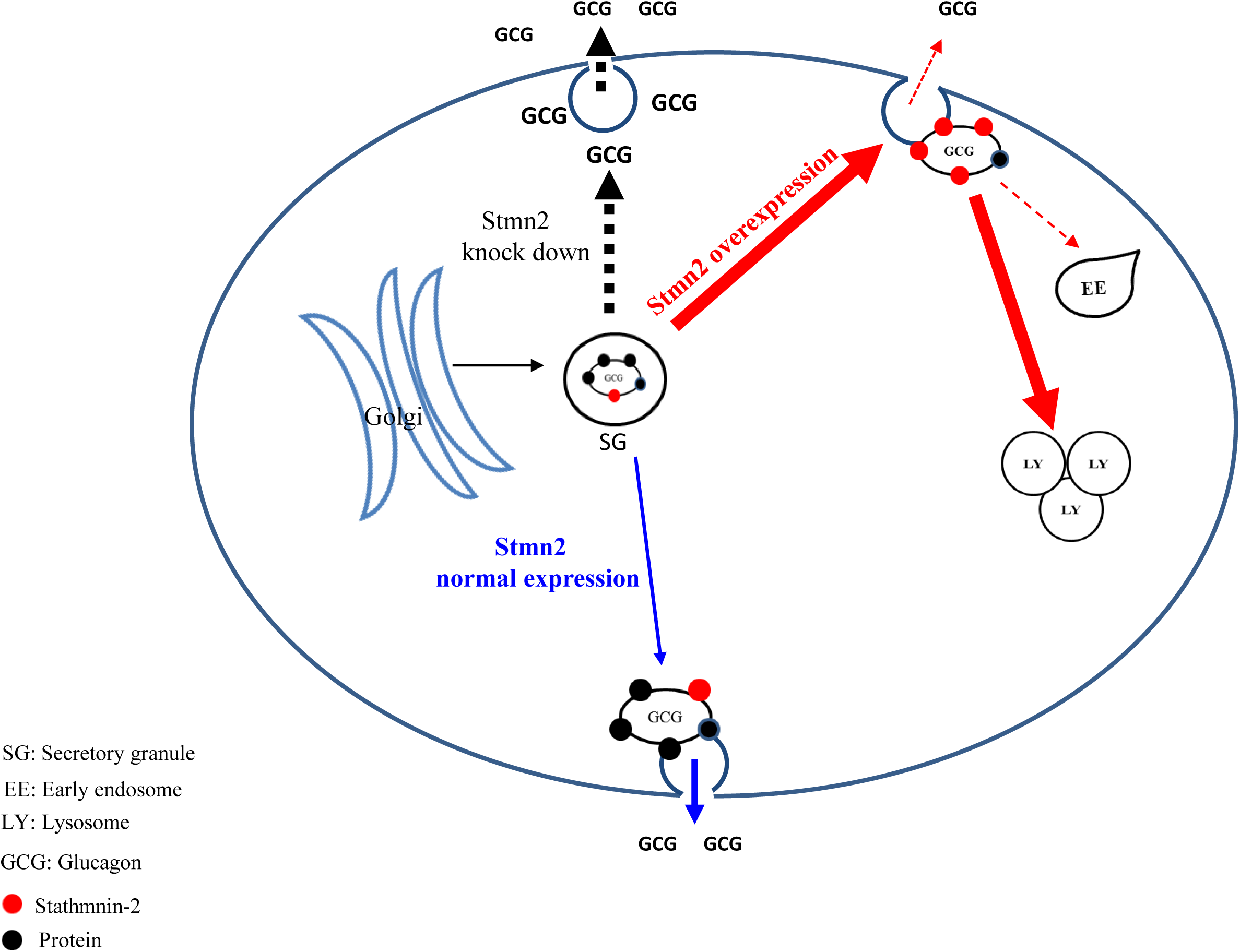
Proposed pathways by which stathmin-2 modulates glucagon secretion from αTC1-6 cells. The model denotes status of glucagon secretion from αTC1-6 cells in normal physiology or in the event of Stmn2-depletion or overexpression. Blue arrows indicate the normal trafficking of glucagon, together with Stmn2, to secretory granules, where they are stored until their release is triggered by a stimulus. Stmn2 overexpression (red arrows) reduces the amount of glucagon available for secretion by diverting secretory granules to lysosomes. Stmn2 depletion (black arrows) reduces the trafficking of glucagon into secretory granules and promotes the constitutive release of glucagon.

## Acknowledgements

We would like to sincerely thank Dr. Jens Juul Holst (Department of Biomedical Sciences, University of Copenhagen, Denmark) for reviewing the manuscript and providing constructive suggestions.

## References

1. Ceriello A, Kilpatrick ES. Glycemic variability: Both sides of the story. Diabetes Care (2013) 36:S272–275. doi:10.2337/dcS13-2030

2. Demant M, Bagger J, Suppli M, Lund A, Gyldenløve M, Hansen K, et al. Determinants of fasting hyperglucagonemia in patients with type 2 diabetes and nondiabetic control subjects. Metab Syndr Relat Disord (2018) 16:530–536. doi:10.1089/met.2018.0066

3. Brand C, Rolin B, Jørgensen P, Svendsen I, Kristensen J, Holst J. Immunoneutralization of endogenous glucagon with monoclonal glucagon antibody normalizes hyperglycaemia in moderately streptozotocin-diabetic rats. Diabetologia (1994) 37:985–993.

4. Gelling RW, Du XQ, Dichmann DS, Romer J, Huang H, Cui L, et al. Lower blood glucose, hyperglucagonemia, and pancreatic alpha cell hyperplasia in glucagon receptor knockout mice. Proc Natl Acad Sci U S A (2003) 100:1438–43. doi:10.1073/pnas.0237106100

5. Holst JJ, Albrechtsen NJW, Pedersen J, Knop FKG. Glucagon and amino acids are linked in a mutual feedback cycle: The liver-α-cell axis. Diabetes (2017) 66:235–240. doi:10.2337/db16-0994

6. Longuet C, Sinclair EM, Maida A, Baggio LL, Charron MJ, Drucker DJ. Response To Fasting. Cell (2009) 8:359–371. doi:10.1016/j.cmet.2008.09.008.The

7. Unger RH, Cherrington AD. Glucagonocentric restructuring of diabetes: A pathophysiologic and therapeutic makeover. J Clin Invest (2012) 122:4–12. doi:10.1172/JCI60016

8. Gromada J, Chabosseau P, Rutter GA. The α-cell in diabetes mellitus. Nat Rev Endocrinol (2018) 14:694–704. doi:10.1038/s41574-018-0097-y

9. Ramracheya R, Chapman C, Chibalina M, Dou H, Miranda C, González A, et al. GLP-1 suppresses glucagon secretion in human pancreatic alpha-cells by inhibition of P/Q-type Ca2+ channels. Physiol Rep (2018) 6:1–17. doi:10.14814/phy2.13852

10. Tornehave D, Kristensen P, Rømer J, Knudsen LB, Heller RS. Expression of the GLP-1 receptor in mouse, rat, and human pancreas. J Histochem Cytochem (2008) 56:841–851. doi:10.1369/jhc.2008.951319

11. Benner C, van der Meulen T, Cacéres E, Tigyi K, Donaldson CJ, Huising MO. The transcriptional landscape of mouse beta cells compared to human beta cells reveals notable species differences in long non-coding RNA and protein-coding gene expression. BMC Genomics (2014) 15:620. doi:10.1186/1471-2164-15-620

12. Ørgaard A, Holst JJ. The role of somatostatin in GLP-1-induced inhibition of glucagon secretion in mice. Diabetologia (2017) 60:1731–1739. doi:10.1007/s00125-017-4315-2

13. Kawamori D, Kulkarni RN. Insulin modulation of glucagon secretion: the role of insulin and other factors in the regulation of glucagon secretion. Islets (2009) 1:276–279. doi:10.4161/isl.1.3.9967

14. Rorsman P, Berggren P, Bokvist K, Ericson H, Möhler H, Ostenson C, et al. Glucose-inhibition of glucagon secretion involves activation of GABAA-receptor chloride channels. Nature (1989) 341:233–236. doi:10.1038/341233a0

15. Basco D, Zhang Q, Salehi A, Tarasov A, Dolci W, Herrera P, et al. α-cell glucokinase suppresses glucose-regulated glucagon secretion. Nat Commun (2018) 9:546. doi:10.1038/s41467-018-03034-0

16. Knudsen J, Hamilton A, Ramracheya R, Tarasov A, Brereton M, Haythorne E, et al. Dysregulation of glucagon secretion by hyperglycemia-induced sodium-dependent reduction of ATP production. Cell Metab (2018)1–13. doi:10.1016/j.cmet.2018.10.003

17. González-Vélez V, Dupont G, Gil A, González A, Quesada I. Model for glucagon secretion by pancreatic α-cells. PLoS One (2012) 7:e32282. doi:10.1371/journal.pone.0032282

18. Wendt A, Birnir B, Buschard K, Gromada J, Salehi A, Sewing S, et al. Glucose inhibition of glucagon secretion from rat α-cells is mediated by GABA released from neighboring β-cells. Diabetes (2004) 53:1038–1045. doi:10.2337/diabetes.53.4.1038

19. Rorsman P, Ashcroft FM. Pancreatic β-Cell electrical activity and insulin secretion: of mice and men. Physiol Rev (2018) 98:117–214. doi:10.1152/physrev.00008.2017

20. Le Marchand SJ, Piston DW. Glucose suppression of glucagon secretion: Metabolic and calcium responses from α-cells in intact mouse pancreatic islets. J Biol Chem (2010) 285:14389–14398. doi:10.1074/jbc.M109.069195

21. Asadi F, Dhanvantari S. Plasticity in the glucagon interactome reveals proteins that regulate glucagon secretion in alpha TC1-6 cells. Front Endocrinol (2019) 9:792. doi:10.3389/fendo.2018.00792

22. Charbaut E, Chauvin S, Enslen H, Zamaroczy S, Sobel A. Two separate motifs cooperate to target stathmin-related proteins to the Golgi complex. J Cell Sci (2005) 118:2313–2323. doi:10.1242/jcs.02349

23. Mahapatra NR, Taupenot L, Courel M, Mahata SK, O’Connor DT. The trans-golgi proteins SCLIP and SCG10 interact with chromogranin A to regulate neuroendocrine secretion. Biochemistry (2008) 47:7167–7178. doi:10.1021/bi7019996

24. Bramswig N, Everett L, Schug J, Dorrell C, Liu C, Luo Y, et al. Epigenomic plasticity enables human pancreatic alpha to beta cell reprogramming. J Clin Invest (2013) 123:1275–1284. doi:10.1172/JCI66514

25. Lawlor N, George J, Bolisetty M, Kursawe R, Sun L, Sivakamasundari V, et al. Single-cell transcriptomes identify human islet cell signatures and reveal cell-type – specific expression changes in type 2 diabetes. Genome Res (2017) 27:208–222. doi:10.1101/gr.212720.116.Freely

26. Mossé Y, Laudenslager M, Longo L, Cole K, Wood A, Attiyeh E, et al. Identification of ALK as the major familial neuroblastoma predisposition gene. Nature (2009) 455:930– 935. doi:10.1038/nature07261.Identification

27. Guizzetti L, McGirr R, Dhanvantari S. Two dipolar alpha-helices within hormone-encoding regions of proglucagon are sorting signals to the regulated secretory pathway. J Biol Chem (2014) 289:14968–14980. doi:10.1074/jbc.M114.563684

28. Zinchuk V, Wu Y, Grossenbacher-Zinchuk O. Bridging the gap between qualitative and quantitative colocalization results in fluorescence microscopy studies. Sci Rep (2013) 3:1– 5. doi:10.1038/srep01365

29. Chauvin S, Sobel A. Neuronal stathmins: A family of phosphoproteins cooperating for neuronal development, plasticity and regeneration. Prog Neurobiol (2015) 126:1–18. doi:10.1016/j.pneurobio.2014.09.002

30. Franklin IK, Wollheim CB. GABA in the endocrine pancreas: its putative role as an islet cell paracrine-signalling molecule. J Gen Physiol (2004) 123:185–190. doi:10.1085/jgp.200409016

31. Daily NJ, Boswell KL, James DJ, Martin TFJ. Novel interactions of CAPS (Ca2+-dependent activator protein for secretion) with the three neuronal SNARE proteins required for vesicle fusion. J Biol Chem (2010) 285:35320–35329. doi:10.1074/jbc.M110.145169

32. Schafer MKH, Mahata SK, Stroth N, Eiden LE, Weihe E. Cellular distribution of chromogranin A in excitatory, inhibitory, aminergic and peptidergic neurons of the rodent central nervous system. Regul Pept (2010) 165:36–44. doi:10.1016/j.regpep.2009.11.021

33. Atouf F, Czernichow P, Scharfmann R. Expression of neuronal traits in pancreatic beta cells. J Biol Chem (1997) 272:1929–1934. doi:10.1074/jbc.272.3.1929

34. Martin D, Grapin-Botton A. The importance of REST for development and function of beta cells. Front Cell Dev Biol (2017) 5:1–13. doi:10.3389/fcell.2017.00012

35. Koshimizu H, Kim T, Cawley N, Loh Y. Chromogranin A: A new proposal for trafficking, processing and induction of granule biogenesis. Regul Pept (2010) 165:95– 101. doi:doi:10.1016/j.regpep.2010.09.006

36. Lutjens R, Igarashi M, Pellier V, Blasey H, Di Paolo G, Ruchti E, et al. Localization and targeting of SCG10 to the trans-Golgi apparatus and growth cone vesicles. Eur J Neurosci (2000) 12:2224–2234. doi:10.1046/j.1460-9568.2000.00112.x

37. Nanjappa V, Thomas J, Marimuthu A, Muthusamy B, Radhakrishnan A, Sharma R, et al. Plasma proteome database as a resource for proteomics research: 2014 update. Nucleic Acids Res (2014) 42:959–965. doi:10.1093/nar/gkt1251

38. Levy A, Devignot V, Fukata Y, Fukata M, Sobel A, Chauvin S. Subcellular Golgi localization of stathmin family proteins is promoted by a specific set of DHHC palmitoyl transferases. Mol Biol Cell (2011) 22:1930–1942. doi:10.1091/mbc.E10-10-0824

39. Pasquier A, Vivot K, Erbs E, Spiegelhalter C, Zhang Z, Aubert V, et al. Lysosomal degradation of newly formed insulin granules contributes to β cell failure in diabetes. Nat Commun (2019) 10:1–14. doi:10.1038/s41467-019-11170-4

40. Yuxia Z, Zhiyu L, Shengmei Z, Ruijuan Z, Huiying L, Xiaoqing L, et al. RILP restricts insulin secretion through mediating lysosomal degradation of proinsulin. Diabetes (2019) doi:10.2337/db19-0086

